# Quantitative MRI-based assessment of the ultrastructural compactness of myelin lamellae

**DOI:** 10.64898/2026.09.30.755666

**Authors:** Mario Corral-Bolaños, Hanna Vila-Merkle, João Lima, Emma L. Thomson, Robin Krüger, Niccolò Peruzzi, Martin Bech, Marco Pizzolato, Tim B. Dyrby

## Abstract

Efficient neuronal communication relying on rapid saltatory conduction depends not only on the amount of myelin wrapping axons but also on myelin compact organisation. Yet, current quantitative MRI myelin imaging methods, such as Myelin Water Imaging (MWI) and quantitative Magnetization Transfer (qMT), are designed to estimate myelin content and are often treated as interchangeable, overlooking myelin organisation. We introduce Structural Myelin Integrity (SMI), a quantitative MRI metric designed to distinguish myelin compact organisation from content. SMI combines the distinct biophysical sensitivities of MWI and qMT to separate the water confined between myelin layers from the total myelin membrane mass. We evaluated SMI in two rodent models of myelin decompaction and one remyelination model, validating our findings with synchrotron small-angle X-ray scattering and electron microscopy. SMI detected myelin layers decompaction despite largely preserved myelin membranes, revealing structural abnormalities that were not explained by myelin content alone. During remyelination, SMI returned closer to healthy values at early timepoints than myelin-content metrics, consistent with the formation of thin-but-compact sheaths. These findings indicate that myelin content and organisation can dissociate and establish SMI as a non-invasive marker of myelin organisation, compatible with existing clinical MRI approaches for studying white matter changes not detected today.

## 1 Introduction

Current neuroimaging studies of white matter suffer from an important blind spot: a heavy reliance on assessing myelin content while overlooking aspects of myelin ultrastructural integrity that are critical for function, such as the preservation and compaction of myelin layers. To ensure fast conduction velocities (CV) enabled through saltatory conduction, white matter axons in the brain are wrapped by myelin, a protein and lipid-rich sheath composed of multiple tightly packed membrane layers known as myelin lamellae^1,2^. Within compact myelin, adjacent lamellae are separated by only a few nanometres and are maintained in close apposition by specialized structural proteins. Importantly, the functional integrity of myelin, i.e., its capability to ensure CV, depends not only on its presence, but also on the preservation of this highly ordered lamellar architecture. Prior work in animal models has associated myelin lamellar separation, hereafter referred to as myelin decompaction, to impaired or blocked conduction velocities even when myelin content is preserved^3–5^. Notably, histopathological studies have identified myelin decompaction as a potential early biomarker of neurodegenerative diseases like multiple sclerosis^6–8^ or Alzheimer’s disease^9^.

Despite its functional implications, myelin lamellar compactness is an under-researched factor in brain health. Currently, electron microscopy (EM) is the only method capable of directly visualizing myelin decompaction, albeit in small fields of view. Moreover, distinguishing pathological decompaction from unspecific myelin degeneration due to tissue fixation artifacts or postmortem times is notoriously difficult^10^. These limitations have hindered the reliable characterization of myelin decompaction across tissues, subjects, and disease populations, and consequently our understanding of how myelin decompaction relates to neurodegenerative disease remains limited. Conversely, magnetic resonance imaging (MRI) enables non-invasive, whole-brain, in vivo imaging. However, current quantitative MRI (qMRI) methods have been designed to estimate the total myelin content voxel-wise rather than assess the structural integrity of the myelin itself^11–15^.

To address the non-invasive assessment of myelin decompaction, we revisited the biophysical underpinnings of two established qMRI techniques and combined them to derive a candidate metric of myelin decompaction. Specifically, we considered Myelin Water Imaging (MWI)^16^, which is sensitive to water confined between myelin lamellae and thus reflects their structural organisation; and quantitative Magnetization Transfer (qMT)^17^, which quantifies the total macromolecular content (mostly lipids) of the myelin sheath in the voxel. Because these techniques probe different physical compartments —fluid spaces (MWI) and structural mass (qMT)— we hypothesised that jointly interpreting these two metrics would be more sensitive to myelin lamellar spacing changes than either metric alone. We reasoned that in healthy, compact myelin, these compartments are expected to maintain a stable relationship, whereas myelin decompaction would decouple them.

Here, we introduce Structural Myelin Integrity (SMI), a metric that combines the water-specific sensitivity of MWI with the macromolecule-specific sensitivity of qMT to approximate the volume fraction of confined water within the myelin sheath (see Methods). We hypothesised that SMI would index myelin compactness differently from either MWI or qMT alone in tissue where myelin content and decompaction dissociate. To test this, we assessed SMI across three rodent models expected to decouple myelin content from compaction: postmortem degeneration and *Plp1*-null mice, both associated with myelin decompaction despite relatively preserved myelin membrane mass, and remyelination following lysophosphatidylcholine (LPC)-induced demyelination, where newly formed sheaths are thin but may already be compact. All experiments were performed on a 7T preclinical scanner using single-slice acquisitions in situ or ex vivo (see Methods). Finally, to confirm the ultrastructural basis of our MRI findings, we compared SMI against standard ultrastructural imaging on the same brains, including electron microscopy and small-angle X-ray scattering computed tomography (SAXS-CT)^18^.

## 2 Results

### 2.1 Longitudinal changes of myelin metrics during postmortem degeneration

After death, unfixed tissue undergoes progressive degeneration. During the early postmortem interval (PMI), myelin microstructure remains largely preserved. However, as PMI extends, myelin progressively degenerates, showing myelin decompaction, splitting and vacuolization^19^. This process provides a model for studying changes in myelin ultrastructural integrity without major changes in myelin content. We hypothesised that myelin layer loosening would differentially affect the two qMRI methods, MWI and qMT, thus providing complementary information and enabling development of SMI. We performed continuous qMRI measurements of unfixed *in situ* rat brains (n = 5). Data were acquired from 2 to 80 hours PMI using a single-slice axial acquisition on a preclinical 7T MRI scanner (Fig. 1a).

**Figure 1.**
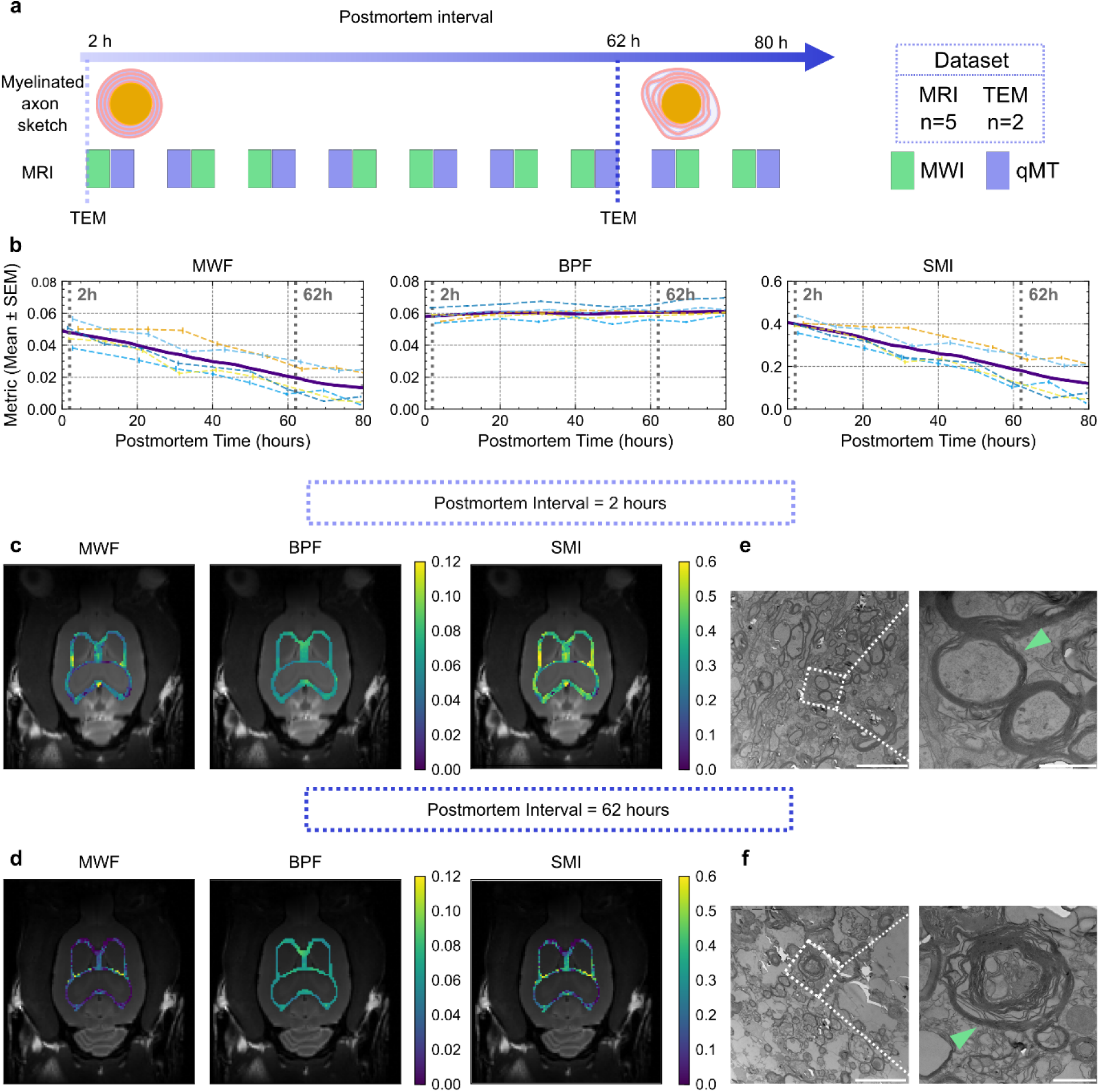
Longitudinal changes of myelin degeneration in unfixed tissue between 2- and 80-hours postmortem. **a.** Experimental design: Five animals were sacrificed and scanned continuously for 80 hours. **b.** Longitudinal changes of the myelin MRI metrics, from left to right: MWF, BPF and SMI. The dashed lines indicate the individual measurements; the purple solid line shows the interpolated average trend through LOWESS regression. **c, d.** Myelin metric maps overlayed in the white matter of a structural image at (**c**) 2 hours and (**d**) 62 hours postmortem. **e, f.** Electron microscope images of the cross-section of white matter axons at (**e**) 2 hours and (**f**) 62 hours postmortem. The green arrows point to the myelin sheath. Scalebar (**e, f**): 5 µm left, 1 µm right.

We observed different trends between the water-based and macromolecule-based metrics as degeneration progressed (Fig. 1b). The Myelin Water Fraction (MWF), which reflects water tightly confined between myelin layers, changed significantly over time (Friedman test: 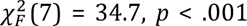) with a negative trend (Page’s *L* trend test: *L* = 1019, n = 5, k = 8, *p* < .001) (Fig. 1b and Supplementary Fig. 1b).

Analysis of the T_2_ spectra revealed that this was driven by a reduction in the myelin water peak amplitude (Supplementary Fig. 1d,e), suggesting that as myelin layers loosen, water escapes the confined short-T_2_ pool. In contrast, the Bound Pool Fraction (BPF), a qMT-derived marker of macromolecular content of the fatty myelin, showed no differences along the degeneration period 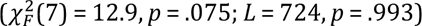 (Fib and Supplementary Fig. 1b).

To assess whether these qMRI trends were accompanied by visible changes in myelin ultrastructure, we examined transmission electron microscopy (TEM) images collected from the corpus callosum of two additional animals (see Methods). At an early timepoint (2 hours PMI), myelin sheaths appeared largely compact, consistent with preserved MWF and BPF (Fig. 1c,e). By 62 hours PMI, however, TEM images showed myelin lamellar decompaction without apparent loss of membrane continuity (Fig. 1f). This specific geometry, myelin decompaction without substantial membrane loss, aligns with our MRI findings. Specifically, the macromolecular content (BPF) remained constant, while the water dynamics within the decompacted layers (MWF) were fundamentally altered (Fig. 1d). Control measures confirmed that MWF decay was not driven by temperature fluctuations, as body temperatures within animals reached room temperature and remained stable within a 2 °C range (Supplementary Fig. 1c). While high body temperatures could bias T_2_ values toward shorter values^20^, thus artifactually reducing MWF, postmortem body temperature progressively declined until stabilizing at room temperature and therefore cannot explain the observed temporal pattern.

To quantify the myelin decompaction effect reflected by the differences between MWF and BPF trends, we formulated the Structural Myelin Integrity (SMI). Mathematically, SMI combines the water-specific sensitivity of MWF with the macromolecule-specific sensitivity of qMT (see Methods). Conceptually, it represents the normalised ratio of the confined water between bilayers relative to the total volume of the myelin sheath (confined water and the macromolecular mass). Since SMI depends positively on MWF at approximately constant BPF, its pattern over the postmortem interval is expected to track the MWF decline shown above. Accordingly, SMI decreased over the same period resembling the trajectory of MWF, consistent with BPF’s stability over the same interval (Fig. 1b), and with the TEM appearance of preserved membrane continuity with lamellar separation (Fig. 1e,f).

### 2.2 Structural validation of SMI in a genetic model of myelin decompaction

Once we established in the postmortem model that the dissociation between MWF and BPF changes reflect myelin decompaction, we next assessed whether SMI is sensitive to genetically induced alterations in myelin architecture using *Plp1*-null mice, a model characterized by widespread myelin decompaction despite largely preserved myelin content^10,21^. We compared the qMRI metrics in a region of interest of the corpus callosum of *Plp1*-null mice (n = 5) versus control mice (n = 4). Then, myelin decompaction on *Plp1*-null mice was validated using SAXS-CT and electron microscopy on the same MRI-scanned mouse brains (Fig. 2a).

**Figure 2.**
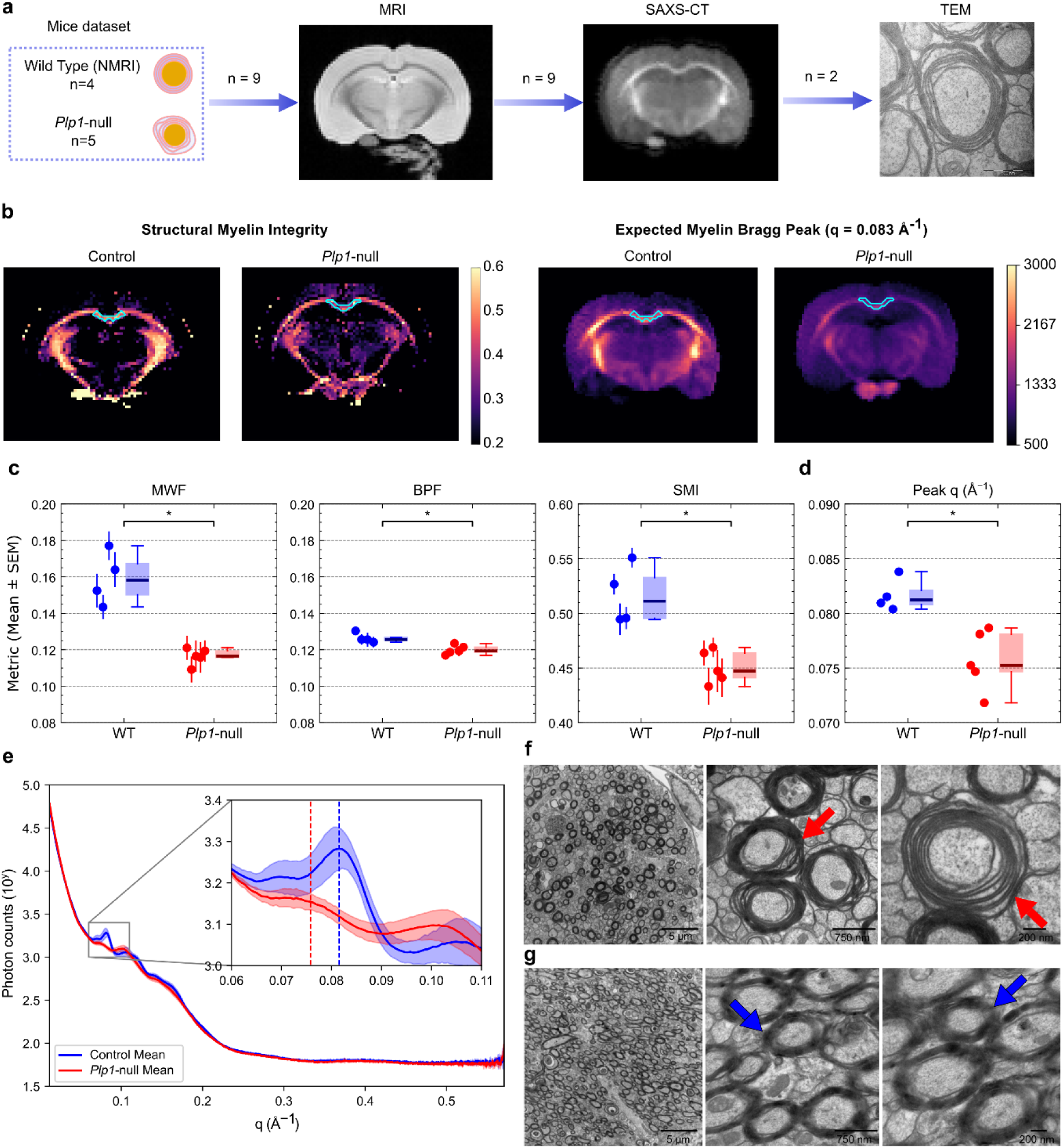
Comparison of qMRI techniques sensitivity to myelin loosening on *Plp1*-null mice. **a.** Experimental design, with the same mouse brains scanned with MRI and SAXS-CT (n=5 *Plp1*-null vs n=4 control mice), and one sample from each group examined with TEM. **b.** Comparison of the derived maps in control mice and *Plp1*-null. In cyan, 20-voxel ROI used for analysis. Left: Structural myelin integrity. Right: SAXS-CT intensity for q values where the myelin peak is expected q = 0.0830 ± 0.0014 Å^-1^. **c,d.** Boxplot of the distribution of the mean metrics in each group. Each point represents the mean ± SEM value per animal in the ROI shown in b. **e.** Mean ± SD scattering intensity in the ROI from each group and estimated myelin peak position (dashed line) against the reciprocal vector q. **f,g.** Transmission electron microscopy images of myelin-stained of the corpus callosum of (**f**) a *Plp1*-null mouse and (**g**) a control mouse. Scalebar (**f, g**): from left to right, 5 µm, 750 nm and 200 nm.

All three qMRI-based myelin metrics —MWF, BPF and SMI— completely separated *Plp1*-null and control animals (Mann–Whitney *U* test: all *U* = 20, *p* = .016) (Fig. 2c). However, the magnitude of the effect differed across metrics. The myelin loosening phenotype drove a 26.3% median decrease in the MWF (Mdn = 0.116, IQR = 0.115–0.119) with respect to controls (Mdn = 0.158, IQR = 0.150–0.167), whereas BPF showed a modest 5.0% reduction (*Plp1*-null: Mdn = 0.119, IQR = 0.119–0.121; controls: Mdn = 0.126, IQR = 0.125–0.127). Thus, the decrease in confined water-associated signal substantially exceeded any change in the macromolecular pool. Consistent with this dissociation, SMI showed a robust 12.5% reduction between *Plp1*-null (Mdn = 0…7, IQR = 0.4411-0.464) compared to controls (Mdn = 0.511, IQR = 0.495–0.533), indicating the impaired structural integrity of the myelin sheath without being confounded by baseline myelin content differences.

To independently confirm the underlying myelin structure changes, we performed whole-brain SAXS-CT, a technique sensitive to the nanometre-scale periodicity of the myelin lamellae, at the same resolution as MRI on the same brains (Fig. 2b)^18^. We observed a significant shift (*U* = 20, *p* = .016) in the Bragg peak toward lower frequencies (*q*-values) in *Plp1*-null mice (Mdn = 0.0752, IQR = 0.07.7–0.0781 Å^-1^) compared to controls (Mdn = 0.0812, IQR = 0.0808–0.0821 Å^-1^) (Fig. 2d). Physically, a lower *q*-value corresponds to a larger *d*-spacing (periodicity), indicating an increase in the periodic spacing between myelin lamellae, consistent with myelin decompaction. Moreover, the averaged scattering profiles showed a reduction in peak intensity (Fig. 2e), suggesting a general loss of periodicity within the myelin sheath.

TEM provided direct visual confirmation of these myelin lamellar alterations and was consistent with previous reports of the *Plp1*-null phenotype (Fig. 2f,g)^22^. Compared with controls, *Plp1*-null mice showed widespread decompaction of myelin lamellae throughout the imaged region (red arrows), while myelin remained compact in controls (blue arrows). Together, SAXS-CT and TEM on the same MRI-scanned brains confirmed extensive myelin decompaction in Plp1-null mice, providing a microstructure explanation for the larger reduction in MWF relative to BPF and the associated decline in SMI.

### 2.3 SMI identifies the compact state of newly formed myelin sheaths during remyelination

To evaluate the sensitivity of SMI to the restoration of compact myelin lamellae following injury, we used LPC to induce a focal demyelinating lesion in the midsagittal corpus callosum of rats. In this model, demyelination peaks at 5–7 days post-injection (dpi), followed by a progressive remyelination^23–26^. We characterized this trajectory in independent groups of animals at 7, 14, 21, and 28 dpi (n=5 per timepoint; Fig. 3a). For each animal, qMRI measurements within the lesion were compared with those from the contralateral control region (Fig. 3b,d). To characterize temporal changes while reducing variability related to lesion location and partial-volume effects associated with the relatively thick acquisition slices, we calculated the lesion-to-contralateral ratio for each qMRI metric (Fig. 3e). We additionally scanned an independent cohort of n=5 naive reference, non-injected rats using the same ex vivo protocol and computed an analogous hemisphere ratio (right-hemisphere ROI over left-hemisphere ROI), providing a lesion-free naive reference distribution for each qMRI metric.

**Figure 3.**
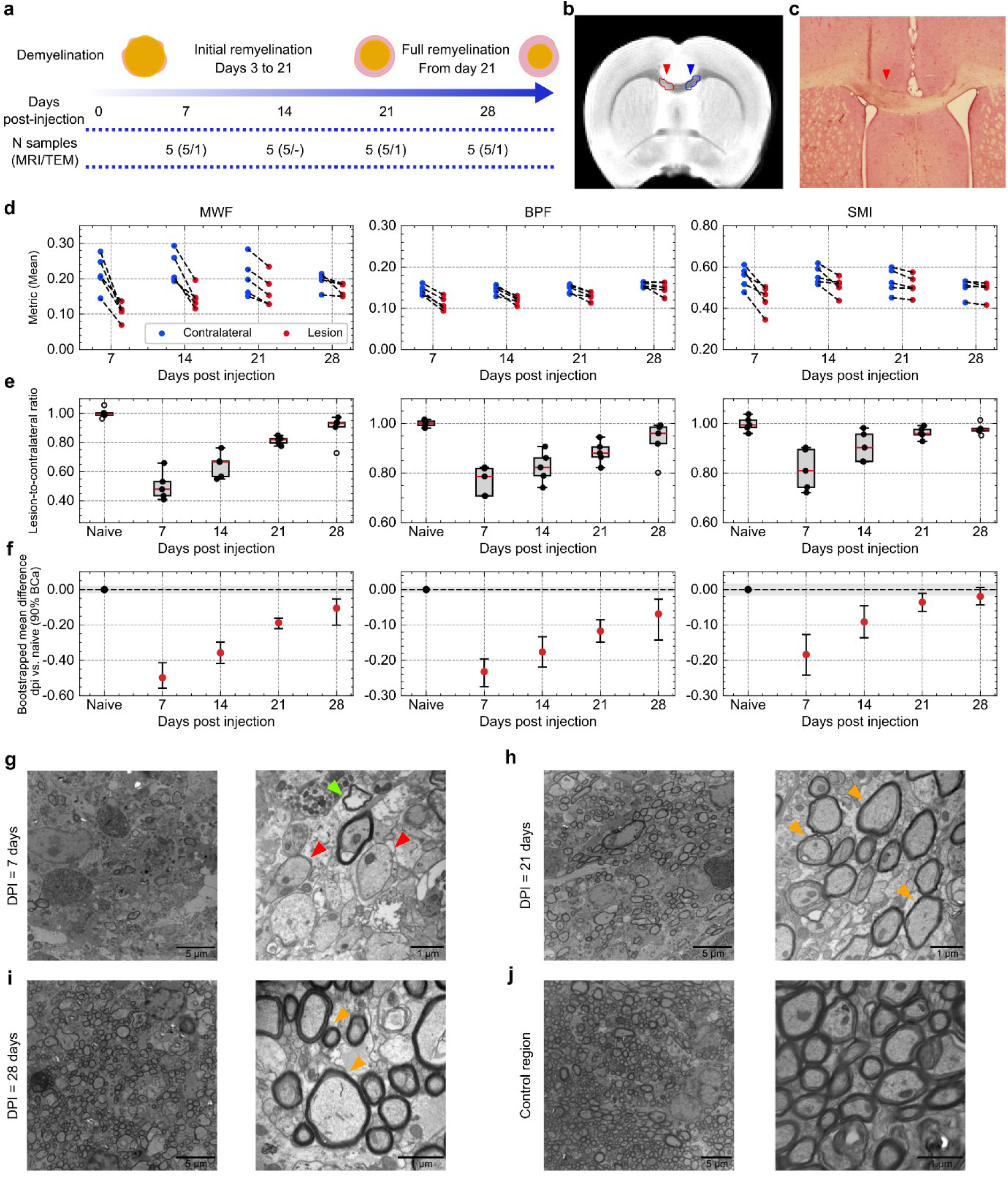
Quantitative MRI study of the remyelination process after LPC-induced demyelination in the rat corpus callosum. **a**. Experimental design. A total of 20 animals were divided in 4 groups depending on which dpi their brains were scanned. Additional 5 non-injected animals were used as naive distribution references. **b**. Representative T_2_w image (dpi=14). In red, the selected region of interest in the lesion area (seen as an hyperintensity). In blue, a contralateral control area. **c**. H&E staining on an LPC-induced lesion (red arrow) on a different rat brain. **d**. Pair-wise comparison of each animal between the lesion and control regions for each metric. **e.** Ratio of the lesion and contralateral area for each qMRI metric across dpi and the naive reference samples. **f**. Estimated bootstrapped mean difference (red dots) ± 90% BCa between each dpi and the naive reference distribution. The shaded area represents the equivalence-margin interval to the reference distribution. **g-j.** Representative TEM images for (**g**) 7 dpi (**h**) 21 dpi (**i**) 28 dpi and (**j**) contralateral control region. Red markers point demyelinated axons, green to myelin debris, and orange to remyelinating axons. Scalebar (**g-j**): 5 µm left, 1 µm right.

At the expected peak of the demyelination (7 dpi), all three metrics showed reduced lesion-to-control ratios. The drop in MWF was the largest (Mdn = 0.48, IQR = 0.43–0.53) compared to a milder reduction in BPF (Mdn = 0.79, IQR = 0.71–0.82). Haematoxylin and eosin (H&E) staining at 7 dpi revealed a marked inflammatory response characterized by increased cellular infiltration (Fig. 3c)^24^. The presence of inflammatory cells, debris, and other non-myelin macromolecules may have contributed to the smaller reduction in BPF, consistent with its known sensitivity to macromolecular content beyond myelin^27–29^. In contrast, the breakdown of the myelin ultrastructure, is expected to reduce the water compartments contributing to MWF. TEM supported the presence of generalized myelin disruption at 7 dpi, showing predominantly demyelinated axons and scattered myelin debris (Fig. 3g). At 7 dpi, SMI also showed a modest reduction relative to the contralateral control region (Mdn = 0.81, IQR = 0.74–0.89), despite the pronounced inflammatory response as observed on H&E staining (Fig. 3c).

During the repair phase (14–28 dpi), lesion-to-control ratios progressively increased toward control values (Fig. 3e). To test recovery formally, we first compared each metric’s lesion-to-contralateral ratio across the reference distribution and the four dpi groups using an omnibus Kruskal-Wallis test; a significant overall difference was observed for all three metrics (MWF: *H*(.) = 21.43, *p* < .001; BPF: *H*(.) = 17.45, *p* = .0016; SMI: *H*(4) = 15.79, *p* = .0034). Subsequently, we assessed recovery between the later timepoints (14, 21 and 28 dpi) and the naive reference distribution through Benjamini–Hochberg adjusted Steel-type tests. MWF remained significantly different from the naive reference distribution at every dpi (14 and 21 dpi, *p*_adj_ = .018; 28 dpi, *p*_adj_ = .024). BPF also remained significantly different at every dpi (14 and 21 dpi, *p*_adj_ = .018; 28 dpi, *p*_adj_ = .0.1). Notably, SMI was only significant at 14 dpi (*p*_adj_ = .02.), but not significantly different at either 21 or 28 dpi (*p*_adj_ = .063 and *p*_adj_ = .22, respectively), the only metric showing this pattern at both timepoints.

To test whether each metric’s ratio was statistically equivalent to the naive reference distribution, rather than simply not significantly different from it, we performed a bootstrap equivalence test against a pre-specified margin (half the standard deviation of the naive reference distribution) at 21 and 28 dpi (Fig. 3f). No metric reached formal equivalence at either timepoints (Table 1). Using the same equivalence framework, a complementary test indicated that no metric’s ratio stabilized between 21 and 28 dpi (i.e., the differences between 21 and 28 dpi were larger than the defined margin) (Table 1). A prospective power analysis showed that formally establishing equivalence or stabilization for the SMI would require 33–43 animals per group, exceeding feasible cohort sizes under this design (Supplementary Note 1). Still, we observed that the estimated differences from naive for MWF and BPF exceeded the pre-specified margin at every tested timepoint, suggesting these differences more plausibly reflect non-negligible deviation from baseline rather than a power limitation alone. Conversely, SMI’s confidence intervals were the only ones to partially overlap the equivalence margin at both timepoints, consistent with a potential near-complete normalization of myelin structural organisation by 21 dpi and subsequent small changes.

**Table 1.**
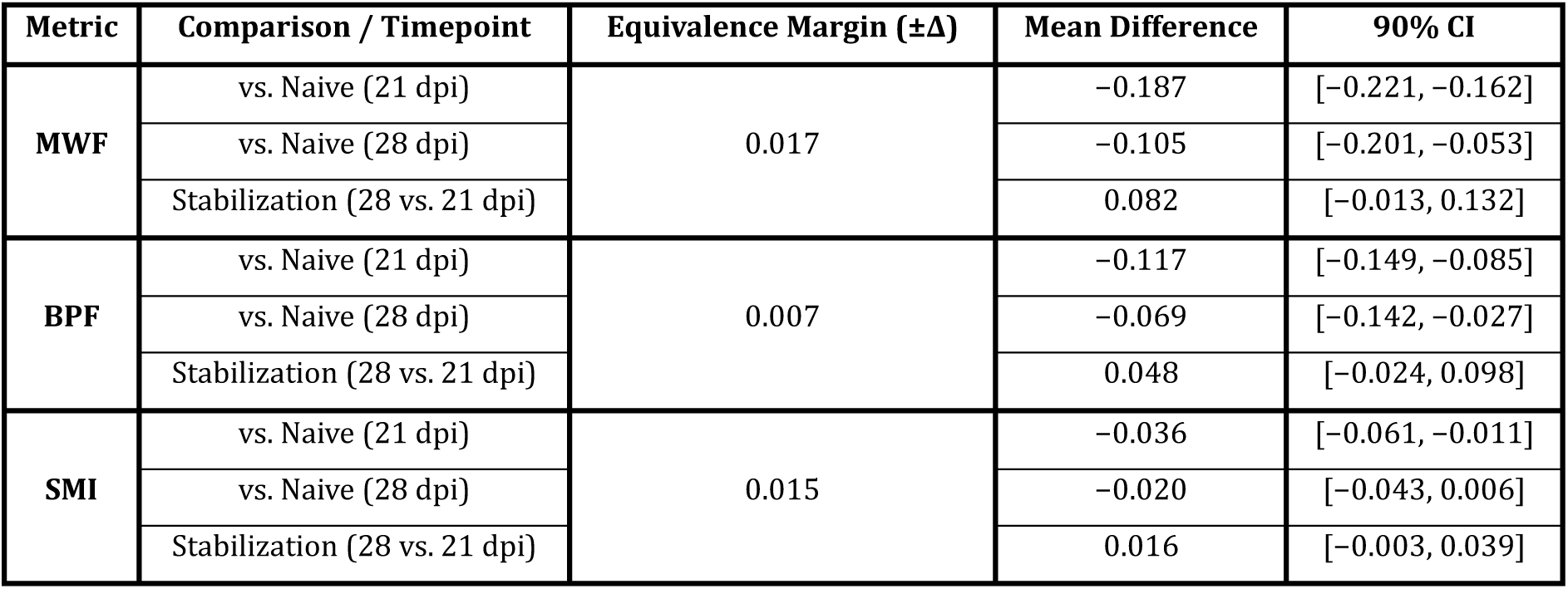
Bootstrap-equivalence tests for baseline recovery and late-stage stabilization. Summary of mean differences, predefined equivalence margins (Δ) and bootstrap 90% confidence intervals (CIs) for MWF, BPF, and SMI compared to naive controls (21 and 28 dpi) and across late timepoints (28 vs. 21 dpi). Equivalence requires the 90% CI to fall entirely within [-Δ, + Δ].

| Metric | Comparison / Timepoint | Equivalence Margin ($\pm\Delta$) | Mean Difference | 90% CI |
| --- | --- | --- | --- | --- |
| MWF | vs. Naive (21 dpi) | 0.017 | -0.187 | [-0.221, -0.162] |
|  | vs. Naive (28 dpi) |  | -0.105 | [-0.201, -0.053] |
|  | Stabilization (28 vs. 21 dpi) |  | 0.082 | [-0.013, 0.132] |
| BPF | vs. Naive (21 dpi) | 0.007 | -0.117 | [-0.149, -0.085] |
|  | vs. Naive (28 dpi) |  | -0.069 | [-0.142, -0.027] |
|  | Stabilization (28 vs. 21 dpi) |  | 0.048 | [-0.024, 0.098] |
| SMI | vs. Naive (21 dpi) | 0.015 | -0.036 | [-0.061, -0.011] |
|  | vs. Naive (28 dpi) |  | -0.020 | [-0.043, 0.006] |
|  | Stabilization (28 vs. 21 dpi) |  | 0.016 | [-0.003, 0.039] |

**Table 2.**
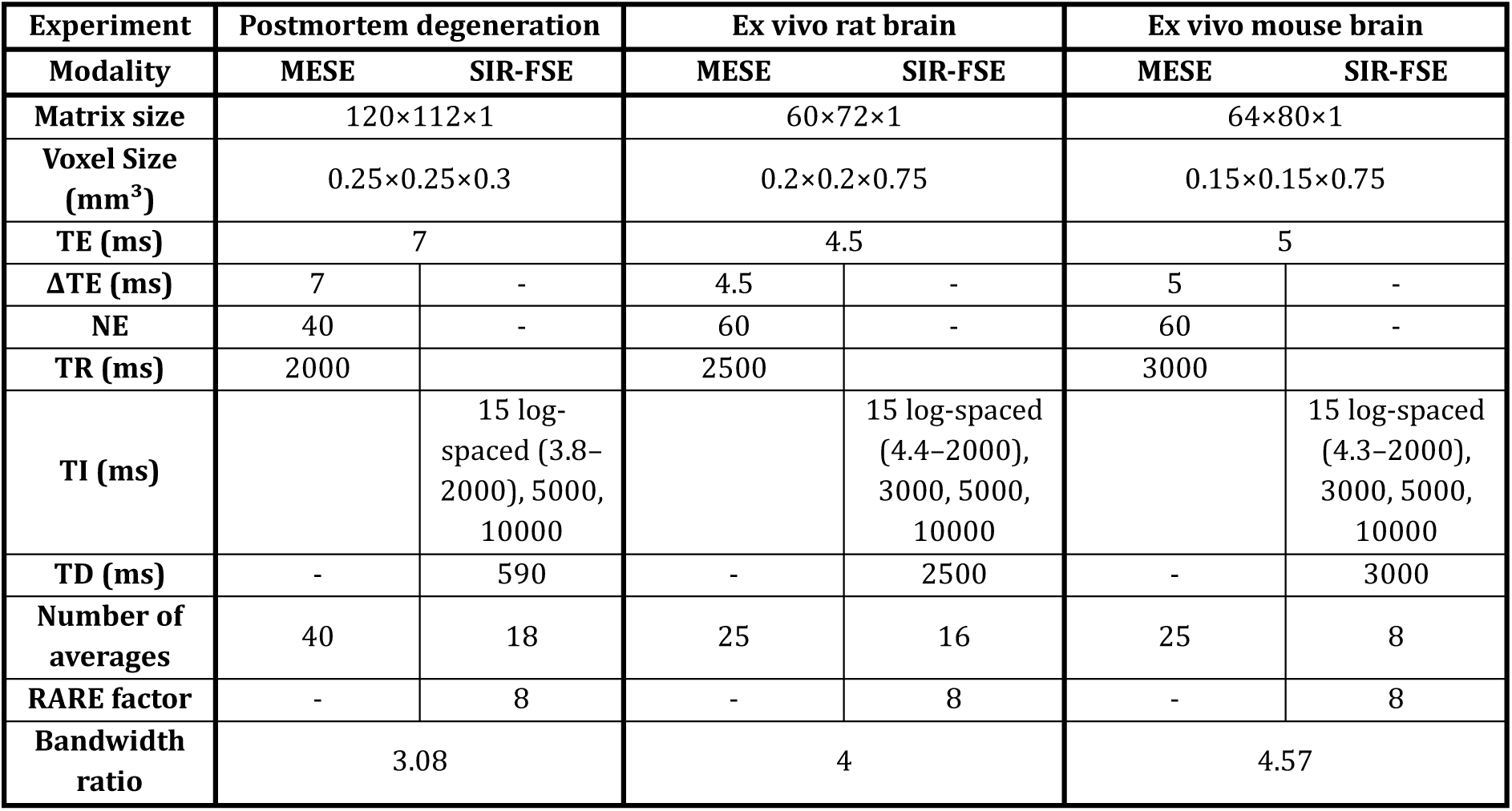
Preclinical acquisition parameters for qMRI experiments. Abbreviations: TE (echo time), ΔTE (spacing between echoes), TR (repetition time), TI (inversion time), TD (delay time).

| Experiment | Postmortem degeneration |  | Ex vivo rat brain |  | Ex vivo mouse brain |  |
| --- | --- | --- | --- | --- | --- | --- |
| Modality | MESE | SIR-FSE | MESE | SIR-FSE | MESE | SIR-FSE |
| Matrix size | 120×112×1 |  | 60×72×1 |  | 64×80×1 |  |
| Voxel Size (mm<sup>3</sup>) | 0.25×0.25×0.3 |  | 0.2×0.2×0.75 |  | 0.15×0.15×0.75 |  |
| TE (ms) | 7 |  | 4.5 |  | 5 |  |
| ΔTE (ms) | 7 | - | 4.5 | - | 5 | - |
| NE | 40 | - | 60 | - | 60 | - |
| TR (ms) | 2000 |  | 2500 |  | 3000 |  |
| TI (ms) |  | 15 log-spaced (3.8–2000), 5000, 10000 |  | 15 log-spaced (4.4–2000), 3000, 5000, 10000 |  | 15 log-spaced (4.3–2000), 3000, 5000, 10000 |
| TD (ms) | - | 590 | - | 2500 | - | 3000 |
| Number of averages | 40 | 18 | 25 | 16 | 25 | 8 |
| RARE factor | - | 8 | - | 8 | - | 8 |
| Bandwidth ratio | 3.08 |  | 4 |  | 4.57 |  |

TEM imaging at 21 and 28 dpi showed remyelination, but the new sheaths were thinner than in contralateral regions, particularly at 21 dpi (Fig. 3h–i). This creates a specific structural phenotype: thin but compact myelin sheaths. Accordingly, the earlier convergence of SMI toward naive reference values is consistent with the hypothesised sensitivity to the organisation of compact myelin lamellar architecture, which appears largely re-established by 21 dpi despite incomplete recovery of myelin thickness. Altogether, our findings suggest that SMI may distinguish the reconstruction of the compacted myelin layers from the accumulation of membrane content.

## 3 Discussion

Myelin water imaging (MWI) and quantitative Magnetization Transfer (qMT) methods have largely been used interchangeably, each one as a proxy of myelin content, with differences between them often attributed to method-specific confounds^28,30^. Our results suggest that those differences carry biological information related to myelin ultrastructure integrity; which, in turn, is known to have a critical influence on the conduction velocities of the nervous impulse^3,5,31^. We assessed three rodent models where myelin content and compactness are expected to dissociate and derived the SMI as an analytical combination of the MWF (MWI-based) and BPF (qMT-based) metrics. The SMI converts the MWF and BPF differences into a quantity with a physical meaning: the fraction of the myelin sheath volume occupied by restricted, short-T_2_ water. Our results indicate that SMI is sensitive to lamellar organisation in a way that neither MWF nor BPF reflect when interpreted alone, and that this sensitivity is consistent across models of ultrastructural myelin alterations.

### 3.1 SMI biophysical interpretation

The performance of SMI across the three animal models is better understood from its compartmental definition (see Methods). Under the assumption that non-myelin macromolecules are negligible, SMI represents the ratio of restricted, short-T_2_ water with respect to the total volume of the myelin sheath. Thus, SMI is a property of the myelin present in the voxel, independent of the total myelin or axonal volume. This opens two scenarios that MWF and BPF cannot distinguish when used individually. First, when myelin lamellae decompact and inter-lamellar restricted water loses its short-T_2_ characterization^32^. In this case, the myelin-water signal declines strongly relative to membrane mass^14^, so MWF decays more than BPF and SMI decreases. The postmortem and *Plp1*-null models correspond to this scenario: MWF was reduced while BPF was relatively preserved, and SAXS-CT and TEM independently showed altered lamellar structure. In our *Plp1*-null experiment, the remaining observed differences in the BPF between groups could possibly reflect baseline myelination differences on the background strain of the mouse models. The second scenario correspond to the case when myelin is thinner but keeps its compact organisation. In this case, the myelin water and myelin macromolecular pools decrease proportionally, so both MWF and BPF are reduced while SMI is preserved. The post-LPC remyelinating experiment shows results consistent with this scenario. First, at 21 and 28 dpi, animals with relatively low MWF and BPF ratios nonetheless showed SMI values close to the naive reference, aligned with the thin but compact sheaths observed using TEM. We additionally observed a consistent reduced variance of SMI at 21 and 28 dpi with respect to its reference naive variance than for the other metrics (Supplementary Note 1). This observation is compatible with the interpretation that late between-animal variance of MWF and BPF could be related to individual biological differences in the myelin reparation process or initial lesion extent^26,33,34^. However, these MWF-BPF shared confounding factors would cancel in SMI, leaving only differences on myelin ultrastructural organisation. Thus, animals at 21 and 28 would then share a largely compacted myelin sheath structure — reflected in the convergence of the SMI distribution— while still differing in how much myelin they have rebuilt —reflected in the remaining variance of MWF and BPF.

This biophysical interpretation of the SMI holds within the assumptions compartmental model, which may be compromised in specific situations. If non-myelin macromolecules contribute substantially to BPF, as during inflammatory infiltration or gliosis (e.g., after LPC injection after 7 days, Fig. 3c), BPF overestimates myelin macromolecular content, and SMI would be biased downwards. In this scenario, SMI would be better considered as a mixed-sensitivity measure than as a specific compaction index and separating myelin from non-myelin macromolecular contributions would require additional constraints, like from inhomogeneous MT^35^. Second, the model assumes negligible water exchange between myelin and non-myelin compartments. However, exchange across thin myelin sheaths has been reported to bias apparent MWF downwards^36,37^. Thus, in newly formed, thin sheaths, the MWF (and hence SMI) could be biased down. In our remyelination experiment, we observed near-normal SMI by 21 dpi, and single animals with low MWF and BPF, but normal SMI (Fig. 3e); but exchange may limit how confidently we can apply the proportional-scaling interpretation for thin-sheaths. In future work, forward simulations of MWI and qMT signals across a range of sheath thickness, lamellar spacings and exchange rates could help to define the influence of these confounds into SMI under different conditions.

### 3.2 SMI among quantitative MRI myelin imaging methods

SMI provides a complementary metric to current quantitative myelin imaging methods. Many established quantitative approaches —such as MT saturation (MTsat)^38^, inhomogeneous MT (ihMT)^39^ or macromolecular tissue volume (MTV)^40^— estimate the macromolecular or myelin lipid content, whereas MWI estimates the volume fraction of water restricted between lamellae^16^. Those methods have typically been validated against histological myelin markers, and comparisons between methods have interpreted their agreement as mutual validation and their differences as method-specific confounding^28,30,41^. Our results suggest a different interpretation. Since MWF depends both on the amount of myelin and the compactness of myelin lamellae, a reduction in MWF cannot differentiate thin myelin sheaths from myelin decompaction^14^. On the other hand, MT-based metrics represent measures of myelin content which are relatively insensitive to myelin ultrastructure. Thus, SMI does not replace either metric; rather, it uses their relationship to differentiate ultrastructural changes of the myelin sheath from changes of content.

The distinction between compact and decompacted myelin has consequences for methods that use myelin imaging as an intermediate quantity. For example, aggregate g-ratio mapping requires an estimate of myelin volume fraction, typically derived from either MWF or MT-based metrics through a calibration factor assumed constant across tissue^42,43^. A fixed calibration between myelin water and myelin volume implicitly assumes a fixed sheath composition, which is equivalent to assuming a constant SMI. Our data indicate that this assumption does not hold in decompacted tissue, where MWF- and MT-based methods to myelin volume fraction would be expected to yield different g-ratio estimates, and more importantly, such estimates would not reflect the actual functional status of the conduction velocities. Beyond its use as an index of compaction, SMI could therefore be used to assess when such calibrations remain valid.

The main cost of SMI is that it requires both MWI and qMT acquisitions. In our preclinical protocol, this came at the expense of single-slice ex vivo acquisitions. However, because SMI is computed analytically from standard MWF and quantitative BPF maps, it requires no additional acquisition in studies that already acquired both and could, in principle, be applied retrospectively to existing datasets, including clinical cohorts. Still, it should be noted that semiquantitative maps, like the ones obtained from MTsat, should be calibrated to represent a macromolecular or myelin volume fraction bounded between 0 and 1^44^. For future clinical studies, faster approaches for estimating MWF and BPF-like maps, such as the single-point method^45^ or accelerated GRASE acquisitions^46^, may further reduce acquisition time.

### 3.3 Towards translation and functional validation

Methods similar methods to the MWI and qMT ones applied in this study have been broadly tested in clinical settings; thus, we expect SMI to be inherently translational. Still, whether applications are informative will depend on the precision with which MWF and BPF can be estimated in vivo. Because SMI combines two independently fitted quantities, its precision depends on both MWF and BPF, and their noise propagation into the ratio. Therefore, translation to clinical imaging will require establishing the SNR at which SMI remains more informative than its inputs; and whether whole-brain MWF and BPF can be acquired together within clinically acceptable scan times.

If these requirements are fulfilled, our results suggest two specific applications. First, the method could be used to assess the presence of myelin decompaction in early stages of neurodegenerative diseases^7,9,47^ or in age-related white matter decline^48,49^. Specifically, histopathological studies have identified myelin decompaction in normal-appearing white matter (NAWM) as an early marker of lesion formation in multiple sclerosis^7,47^. In these conditions, a decrease in SMI should precede, or exceed, a decrease in BPF, meaning that longitudinal imaging could reveal early ultrastructural changes before gross demyelination is detectable by MT-based methods. Moreover, SMI would enable the establishment of fixed diagnostic thresholds. Such a threshold could allow for the identification of “decompacted but present” myelin as an early therapeutic target. Beyond neurodegeneration, SMI could also provide new insight into healthy aging, as myelin undergoes structural changes across the lifespan that are not necessarily reflected in measures of myelin content^48,49^. Similarly, alterations of myelin have been hypothesised to contribute to the pathophysiology of psychiatric disorders^50,51^. In these conditions, SMI could help determine whether white matter differences reflect altered myelin organisation rather than a change in the amount of myelin. Second, in remyelination trials, MWF and MT-based metrics recover slowly and often incompletely because newly remyelinating sheaths usually remain thinner and shorter^52^. A measure that is sensitive to sheath composition rather than amount could indicate structural repair earlier than content-based measures. Importantly, future clinical studies should consider the possible influence of iron content changes since SMI inherits MWF’s sensitivity to tissue iron. For example, iron reduction due to oligodendrocyte loss, could bias MWF downwards changing SMI regardless of myelin lamellar organisation^53,54^. Therefore, applications in scenarios with possible iron content changes should interpret SMI carefully. Potentially, quantitative susceptibility mapping (QSM) could be integrated to disentangle iron content biases from actual myelin compactness.

Finally, the functional significance of the changes detected by SMI remains to be established. While different computational^5^ and experimental^3,55^ studies have shown the importance of myelin ultrastructure in conduction velocities, the exact relationship between conduction velocities, myelin content metrics and SMI remains to be determined. Delays in conduction velocities have been identified in the case of myelin layers decompaction^3,5^, while LPC-induced demyelination models have shown an initial total block of the nervous impulse during demyelination, with a partial recovery from 1. dpi onwards^55^. Future studies should establish direct comparisons between SMI and electrophysiological conduction velocities within the same experimental groups. Such an experiment would establish whether SMI sensitivity to compaction also reflects sensitivity to function and potentially provide a new tool to characterize in vivo and non-invasively the structure-function relationship between myelin and conduction velocities.

## 4 Methods

### 4.1 Animal tissue specimens

All animals in this study were housed under a 12-hour light/dark cycle with ad libitum access to food and water. A total of n=32 female Sprague Dawley rats (NTac:SD), aged 10-16 weeks were used. Of those, 5 were used in the postmortem degeneration study, 2 were used for postmortem TEM imaging, 20 underwent surgery for the LPC-induced demyelination study, and 5 were used as a naive, non-injected reference cohort. A total of n=5 male PLP-null (B6;129-*Plp1^tm1Kan^*/J) (RRID: IMSR_JAX:003255)^56^; and n=. male NMRI mice (BomTac:NMRI), aged 10-12 weeks, were used in this study. Before imaging, all animals were euthanized with an overdose of pentobarbital, and those which were perfused-fixated for ex vivo MRI scanning were transcardially perfused with 4% formaldehyde in neutral buffer (CellPath, UK).

All experimental procedures were approved in advance by the Danish authority responsible for overseeing animal experiments (The Animal Experiments Inspectorate of Denmark, Protocol number 2023-15-0201-01494) and were carried out in compliance with national and international regulations on the care and use of laboratory animals (Directive 2010/63/EU of the European Parliament).

### 4.2 LPC injections

A focal demyelinating lesion in the corpus callosum was induced by an intracranial injection of 1 µl of 1% lysophosphatidylcholine (LPC; Cat. No. L4129, Sigma-Aldrich, St. Louis, MO, USA). LPC primarily disrupts oligodendrocyte membranes, causing cell death and a subsequent inflammatory response that leads to focal demyelination. The LPC intracranial injections were carried out under general anaesthesia with isoflurane, delivered via a calibrated vaporizer (4–5% for induction; 1–2% for maintenance). Each animal was then secured in a stereotaxic apparatus, and key physiological parameters (body temperature, heart rate, and oxygen saturation) were continuously monitored to ensure stability. Under aseptic conditions, the scalp was disinfected, and local anaesthesia was administered subcutaneously. A single midline incision exposed the skull, and coordinates for the corpus callosum (AP: 0.96, DV: −400, ML: 1.00 mm relative to bregma) were determined using a standard stereotaxic atlas^57^. A burr hole was then drilled at these positions using a precision microdrill. A microinjection needle was gently lowered into the target region, where 1 µl of 1% LPC was infused at 0.1 µl/min to minimize mechanical injury. After the injection, the needle remained in place for approximately 10 minutes to reduce the risk of backflow. The incision was closed with absorbable sutures, and the animals were placed in a temperature-controlled chamber until they recovered normal reflexes and mobility.

### 4.3 Preclinical magnetic resonance imaging

All imaging was performed on a 7T horizontal bore Bruker BioSpec 70/20 scanner. For the postmortem degeneration analysis, we used an 86 mm inner diameter Bruker volume coil for transmission and a 20 mm surface coil for reception. For ex vivo acquisitions, we used a 35 mm inner-diameter Bruker quadrature bird-cage coil for transmission and reception. Myelin Water Imaging (MWI) data was acquired through single slice multi-echo spin echo (MESE) sequence; and for quantitative Magnetization Transfer (qMT) imaging we used a single-slice selective inversion-recovery prepared fast spin echo sequence (SIR-FSE)^58^. Acquisition parameters are summarized in Table.

#### Longitudinal postmortem tissue degeneration

Approximately 2 hours after sacrificing, we acquired consecutive blocks of 4-hour MRI scanning protocols in situ during approximately 80 hours. In each block, we acquired and altered the order of MWI and qMT. Body temperature was monitored during the acquisition through a rectal probe.

#### Ex vivo imaging

All ex vivo imaging was performed through single coronal slice acquisitions. Slices were oriented such that they minimized partial volume effects through the corpus callosum. At least 10-days before scanning, brains were placed in 0.1 M PBS (mice brains 10–1. days, rat brains 10–30 days). To prevent susceptibility-induced distortions, ex vivo brains were immersed in a perfluorocarbon solution (Fluorinert^®^) during scanning.

### 4.4 SAXS-CT experiments and reconstruction

The NMRI and *Plp1*-null mouse brains were scanned at the ForMAX beamline^59^ at MAX IV, Lund, Sweden. Tissue preparation for SAXS-CT used the same setup as used in MRI. Postmortem brain tissue stored in PBS was mounted in an Eppendorf tube with cross-sectional dimensions matching the sample, as used for MRI. First, the tube was filled with PBS, and an agar block was pushed to the bottom of the tube. The length of the agar block positioned the brain in the centre of the tube for SAXS-CT imaging. Then, a second agar block was inserted until the tube was filled, securing mechanical stability during physical rotation in the SAXS-CT experiment. Sample preparation was completed at least one day before SAXS-CT measurements, and the mounted sample was kept at room temperature to ensure mechanical stability.

At the ForMAX beamline the undulator emitted X-ray beam was monochromatized using a double crystal monochromator to 20.1 keV and focused using bendable mirrors to a spot size of 50 µm × 50 µm. The samples were mounted in this focus position on top of a stack of stages. Two translation stages, oriented transversal to the beam, enabled to map the sample and create sample projection images. An additional rotation axis enabled to change the projection angle to acquire maps from different directions. The sample was mounted on a goniometer head, on top of the scanning stages, to align the sample manually on to the rotation axis and correct the tilt.

Scattering patterns were collected for each point of the projection maps using an Eiger2 X 4M (Si) photon counting detector located 2.49 m downstream of the sample. In addition to to the scattering signal the transmitted beam was measured using a diode on the beam stop. One SAXS-CT acquisition of a full brain consisted of 40 projection maps with a width of 12 mm including the outer tube diameter and different heights based on the individual sample dimension. A bi-directional step size of 150 µm was used with an exposure time of 30 ms per point. The scan speed was limited by the utilized stages leading to a total scan time per brain of approximately 4 h and 30 min.

An azimuthal integration of the scattering pattern was performed into 1000 q-bins covering a range from 0.0005 Å^-1^to 0.57 Å^-1^. The centre-of-rotation was manually found for each sample using a set of reconstructions of the transmission (diode) data. Using that information the full volumes for transmission and scattering data were reconstructed using the algebraic reconstruction technique (ART) as implemented in the TomoPy toolbox^60^. The scattering data was normalized with the transmission data (diode) prior reconstruction to correct for sample absorption. Each q-bin was reconstructed separately resulting in a full scattering curve for each voxel in the reconstruction volume.

### 4.5 Histological preparation and TEM imaging

#### Epon embedding of fixed tissue

After MRI-imaging, rat (n=3) and mice (n=2) brains were prepared for TEM imaging. For mice, a mid-sagittal slice from the right hemisphere was punctured using a 1 mm diameter biopsy puncture tool, covering the posterior part of the corpus callosum, approximately corresponding to the imaged region with MRI. In the case of rat brains, two different punctures, corresponding to the LPC injection (right hemisphere), and control region on the left hemisphere were taken. Afterwards, the samples were transferred to a 2.5% solution of glutaraldehyde in cacodylate buffer (0.1 M) and stored for two weeks. Then, Epon embedding was conducted following a standard procedure^61^.

#### Epon embedding after postmortem degeneration

Following euthanasia, two brains were carefully extracted and stored at room temperature in phosphate-buffered saline (PBS) for 2 and 62 hours. For electron microscopy, small cross-sectional blocks containing the corpus callosum were dissected and fixed postmortem in 1% osmium tetroxide in 0.1M phosphate buffer (pH 7.4) for 1–2 hours at room temperature. After fixation, the tissue samples were dehydrated in a graded ethanol series (30–100%), followed by infiltration with propylene oxide. The samples were then embedded in Epon resin, polymerized at 60°C for 48 hours, and ultrathin sections (70–90 nm) were cut using an ultramicrotome.

#### TEM imaging

Sections were collected on copper grids, stained with uranyl acetate and lead citrate, and examined on a CM100 microscope (Philips; FEI, Eindhoven, NL). Images were acquired digitally for subsequent visualization.

#### Haematoxylin and eosin (H&E) staining

An LPC-injected rat brain was coronally sectioned at 50 μm using a vibrating blade microtome (VT1000 S, Leica Biosystems, Germany). Tissue sections were mounted on glass slides (Starfrost, Hounisen, Denmark) and air-dried before staining with H&E^24,62^. Staining was performed as follows: immersion in haematoxylin (. min), rinsing under running deionised water (10 min), and immersion in eosin (1 min). Afterwards, the sections were dipped 1–2 times in deionised water, dehydrated with 2–3 dips in 70% alcohol, immersed in 96% alcohol (2 min) and 99% alcohol (two 2.5-min changes), and then cleared in xylene (two 2.5-min changes). Finally, the sections were coverslipped with Tissue-Tek® Glas Pertex (Sakura, Japan) before acquisition of a digital image.

### 4.6 Data analysis

#### Quantitative MRI Myelin Structural Integrity model

Quantitative MRI analysis data was fitted on voxel-wise basis. For qMT analysis, the images were fitted to the Bloch-McConnel equations describing longitudinal relaxation and magnetization transfer between water and macromolecular protons, resulting in estimations of the BPF ^58^. For MWI, we estimated the T_2_ spectrum through a minimum-curvature regularized (λ=5×10^-4^) algorithm. MWF maps were generated with a T_2_ cutoff of 17 ms for unfixed-postmortem tissue and 30 ms for ex vivo tissue analysis.

For the Structural Myelin Integrity (SMI) model, we adopted the four-pool compartmental model described by West et al^28^. We assume the white matter voxel comprises four distinct proton pools: myelin water (V_W,M_), myelin bound protons (V_B,M_), non-myelin water (V_W,NM_), and non-myelin bound protons (V_B,NM_). We assume exchange of longitudinal magnetization between the bound and water protons, but negligible exchange of magnetization or water between myelin and non-myelin compartments. Assuming equal proton molar concentrations across compartments, magnetization fractions are equivalent to volume fractions. Hence, standard quantitative metrics are defined as follows. The Myelin Water Fraction (MWF) represents the fraction of total water localized within the myelin sheath:

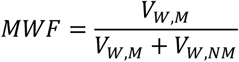

The Bound Pool Fraction (BPF) represents the ratio of all macromolecular-bound protons to the total proton pool:

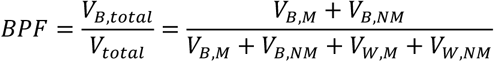

Alternatively, the water volume fraction (WVF) in the voxel is the complementary to the BPF:

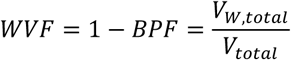

We define the Structural Myelin Integrity (SMI) index as the fraction of myelin water volume with respect to the total myelin volume, i.e., the solid (bound) plus the myelin water volume.

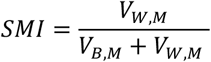

To operationalize this using MRI observables, we assume that myelin is the dominant source of macromolecules in white matter, such that V_B,NM_ ≪ V_B,M_. Under this approximation (V_B,total_ ≈ V_B,M_), the SMI can be derived analytically from the standard MWF and BPF maps:

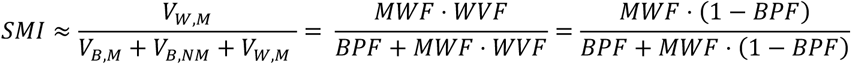

Thus, SMI provides a dimensionless ratio between 0 and 1 of sheath compactness, independent of the total axonal density or total water content in the voxel.

#### SAXS-CT myelin isolation

To isolate the myelin-specific signal in each scattering pattern, we followed the procedure proposed by Georgiadis et al^63^. First, we modelled the background signal by fitting an exponential function, 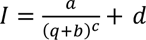, to the q-ranges: q < 0.0. Å^-1^ and q > 0.2 Å^-1^. After subtracting this baseline, residual background was removed by applying a fourth-degree polynomial fit within specific *q*-ranges: for NMRI mice, 0.0.8–0.065, 0.091–0.09., 0.130–0.135, and 0.175–0.210 Å^-1^; for *Plp1*-null mice, 0.065–0.068, 0.086–0.090, 0.130–0.135, and 0.175–0.210 Å^-1^. These ranges differed between NMRI and *Plp1*-null mice to adjust for variations in myelin peak position. Finally, the Bragg q-peak position was identified in the corrected signal within the range 0.068–0.092 A ^−1^ using the scipy.signal.find_peaks function.

### 4.7 Statistical tests

Statistical analyses were performed using open-source Python (v3.12.13) packages SciPy (v1.15.3)^64^, statsmodels (v0.1.46)^65^, and scikit-posthocs (v0.17.0)^66^.

Nonparametric tests were used as distributional assumptions were not considered appropriate. To assess differences across multiple repeated measures or longitudinal timepoints within the same subjects, the Friedman test was used; an ordered directional trend across these sequential datapoints was evaluated using Page’s L trend test. For independent two-group comparisons (e.g., controls vs. Plp1-null mice), the Mann-Whitney *U* test was applied. For the independent many-group comparison (LPC time-course experiment), the Kruskal-Wallis test was implemented, followed by post-hoc Steel-type tests with Benjamini-Hochberg correction. Unless otherwise stated, all statistical tests were two-tailed, and a value of *p* < 405 was considered statistically significant. Locally weighted scatterplot smoothing (LOWESS) regression was utilized to model and visualize the continuous average trend of data collected across varying timepoints. Exact sample sizes (n) for each experiment are indicated in the respective results text and figure legends.

#### Longitudinal postmortem tissue degeneration

Regions of interest covering the white matter (corpus callosum) in axial slices were delineated and metrics were reported as averages within the ROI. To standardize the irregular time-series sampling across different subjects prior to statistical analysis, each metric’s longitudinal profile was linearly resampled at 10-hour intervals spanning 5 to 75 hours postmortem. A LOWESS regression (fraction of neighbours = 0.4) was then used to fit the overall trend. Longitudinal differences across all standardized timepoints were evaluated using the Friedman test, while the presence of a hypothesised monotonic decreasing trend was assessed via Page’s L trend test.

#### Ex vivo mouse brain: *Plp1*-null mice study

Regions of interest of 20 voxels were delineated in the centre of coronal slices in the corpus callosum to minimize partial volume effects. SAXS-CT and qMRI data were aligned to a common space through a rigid registration using elastix^67^. Comparison between groups was performed through Mann-Whitney U tests.

#### Ex vivo rat brain: LPC-induced demyelination and remyelination study

Before analysis, images were interpolated to 0.1×0.1×0.75 voxel size to mitigate partial-volume effects through a cubic B-splines interpolation. Regions of interest of 45 voxels were delineated in the lesion area and a contralateral control region in the opposite hemisphere. For each animal and qMRI metric, the lesion-to-control ratio was calculated to reduce inter-animal variability related to lesion location and partial-volume effects associated with the relatively thick acquisition slices. Additionally, an independent cohort of n=5 naive, non-injected rats was scanned using the same ex vivo protocol, and an analogous hemisphere ratio (right-hemisphere ROI over left-hemisphere ROI) was computed for each metric to provide a lesion-free reference distribution. A ratio of 1 indicated equal values in the lesion and contralateral regions, whereas ratios below 1 indicated lower values in the lesion.

For each metric, an omnibus Kruskal-Wallis test was first performed across the naive reference distribution and the four dpi groups. Then prespecified planned comparisons per metric in the later timepoints (naive vs. 1., 21, 28 dpi) were then performed using two-sided Steel-type tests. *P*-values from the resulting nine comparisons (three metrics times three timepoints) were jointly adjusted using the Benjamini-Hochberg procedure.

To test whether each metric’s lesion-to-contralateral ratio was statistically equivalent to the naive reference distribution, a bootstrap equivalence test was performed at 21 and 28 dpi. For each metric, a pre-specified equivalence margin was defined as half the standard deviation of the naive reference distribution. A 90% bias-corrected and accelerated (BCa) bootstrap confidence interval (10000 resamples) for the mean difference between each dpi group and the naive reference distribution was compared against this margin. Equivalence was accepted if the confidence interval fell within the margin. The same bootstrap-equivalence framework tested whether each metric’s ratio had stabilized between 21 and 28 dpi. To contextualise the null equivalence results, a prospective power analysis was performed to estimate the sample size that would be required for the same equivalence tests to reach 80% power at the pre-specified margin. Required sample sizes were computed analytically using the Welch-corrected two one-sided test (TOST) framework with Satterthwaite approximate degrees of freedom^68^ (further details in Supplementary Note 1).

## Data availability

The authors declare that the source data supporting the findings of this study are available within the paper. Source data for Figures 1, 2, 3, Table 1 and Supplementary Information are provided with the paper.

## Code availability

Quantitative Magnetization Transfer and Myelin Water Fraction data was fitted using the open-source REMMI toolbox (https://github.com/remmi-toolbox/remmi-matlab).

## Competing interests

Authors declare no competing interests.

## Acknowledgements

The project has received funding from the European Research Council (ERC) under the European Union’s Horizon Europe research and innovation programme (grant agreement No. 1010.4180; Principal investigator: Tim B. Dyrby). This work received funding from the Independent Research Fund Denmark (case number 10.465.0/3105-00129B; Principal investigator: MP). We acknowledge the Core Facility for Integrated Microscopy, Faculty of Health and Medical Sciences, University of Copenhagen, for the support in sample preparation and electron microscope imaging. We are thankful for the laboratory assistance of Vibeke Høgsberg in sample preparation for H&E staining and imaging, Centre for Neuroscience and Stereology, Department of Neurology, Bispebjerg and Frederiksberg Hospital. We acknowledge the MAX IV Laboratory for beamtime on the ForMAX beamline under proposal 20250958. Research conducted at MAX IV, a Swedish national user facility, is supported by Vetenskapsra det (Swedish Research Council, VR) under contract 2018-07152, Vinnova (Swedish Governmental Agency for Innovation Systems) under contract 2018-0.969 and Formas under contract 2019-02.96. R.K., N.P. and M.B. acknowledge funding from the European Research Council (ERC) under the European Union’s Horizon 2020 research and innovation program (Grant agreement No. 10108933.). E.T. acknowledges funding from the Novo Nordisk Foundation (grant number NNF00777698, “XtremeCT”).

## Author contributions

T.B.D. conceived the study and coordinated the experiments. M.C.B. acquired and processed the MRI data. M.C.B., M.P. and T.B.D. derived the Structural Myelin Integrity model. H.V.M. and J.L. performed the LPC-surgeries and prepared the ex vivo samples. E.T., R.K., M.B. and T.B.D. acquired the SAXS-CT data. R.K., N.P., and M.B. reconstructed the SAXS-CT data. E.T. and M.C.B. analysed the SAXS-CT data. M.C.B. acquired the TEM data. J.L. performed the H&E histology. M.C.B. and J.L. performed the statistical analysis. M.C.B. wrote the original draft with inputs from T.B.D and M.P. Research was supervised by T.B.D. The text was reviewed and edited with input from all the coauthors.

## Notes

### Competing Interest Statement

The authors have declared no competing interest.

## References

1. Bakiri, Y., Kàradóttir, R., Cossell, L. & Attwell, D. Morphological and electrical properties of oligodendrocytes in the white matter of the corpus callosum and cerebellum. Journal of Physiology 589, 559–573 (2011).

2. Basu, K., Appukuttan, S., Manchanda, R. & Sik, A. Difference in axon diameter and myelin thickness between excitatory and inhibitory callosally projecting axons in mice. Cerebral Cortex 33,4101–4115 (2023).

3. Gutie rrez, R., Boison, D., Heinemann, U. & Stoffel, W. Decompaction of CNS myelin leads to a reduction of the conduction velocity of action potentials in optic nerve. Neurosci. Lett. 195, 93–96 (1995).

4. Rodionova, N. N., Allakhverdiev, E. S. & Maksimov, G. V. Study of myelin structure changes during the nerve fibers demyelination. PLoS One 12, e0185170 (2017).

5. Jamann, N. et al. Layer 5 myelination gates corticothalamic coincidence detection. Nat. Commun. 16, 10922 (2025).

6. van den Bosch, A. M. R., et al. Ultrastructural Axon–Myelin Unit Alterations in Multiple Sclerosis Correlate with Inflammation. Ann. Neurol. 93, 856–870 (2023).

7. Shaharabani, R. et al. Structural Transition in Myelin Membrane as Initiator of Multiple Sclerosis. J. Am. Chem. Soc. 138, 12159–12165 (2016).

8. Bando, Y. et al. Abnormal morphology of myelin and axon pathology in murine models of multiple sclerosis. Neurochem. Int. 81, 16–27 (2015).

9. Depp, C. et al. Myelin dysfunction drives amyloid-β deposition in models of Alzheimer’s disease. Nature 618, 349–357 (2023).

10. Möbius, W., Nave, K. A. & Werner, H. B. Electron microscopy of myelin: Structure preservation by high-pressure freezing. Brain Res. 1641, 92–100 (2016).

11. Schmierer, K. et al. Quantitative magnetization transfer imaging in postmortem multiple sclerosis brain. Journal of Magnetic Resonance Imaging 26, 41–51 (2007).

12. Laule, C. & Moore, G. R. W. Myelin water imaging to detect demyelination and remyelination and its validation in pathology. Brain Pathology 28, 750–76. (2018).

13. McCreary, C. R. et al. Multiexponential T2 and magnetization transfer MRI of demyelination and remyelination in murine spinal cord. Neuroimage 45, 1173–1182 (2009).

14. Chen, H. S.-M., Holmes, N., Liu, J., Tetzlaff, W. & Kozlowski, P. Validating myelin water imaging with transmission electron microscopy in a rat spinal cord injury model. Neuroimage 153, 122–130 (2017).

15. Thiessen, J. D. et al. Quantitative MRI and ultrastructural examination of the cuprizone mouse model of demyelination. NMR Biomed. 26, 1562–1581 (2013).

16. Mackay, A. et al. In vivo visualization of myelin water in brain by magnetic resonance. Magn. Reson. Med. 31, 673–677 (1994).

17. Sled, J. G. & Pike, G. B. Quantitative imaging of magnetization transfer exchange and relaxation properties in vivo using MRI. Magn. Reson. Med. 46, 923–931 (2001).

18. Jensen, T. H. et al. Molecular X-ray computed tomography of myelin in a rat brain. Neuroimage 57, 124–129 (2011).

19. Krassner, M. M. et al. Postmortem changes in brain cell structure: a review. Free Neuropathol. 4, 10 (2023).

20. Rieke, V. & Butts Pauly, K. MR thermometry. Journal of Magnetic Resonance Imaging 27, 376–390 (2008).

21. Rosenbluth, J., Nave, K., Mierzwa, A. & Schiff, R. Subtle myelin defects in PLP-null mice. Glia 54, 172–182 (2006).

22. Groh, J. et al. Pathogenic inflammation in the CNS of mice carrying human PLP1 mutations. Hum. Mol. Genet. 25, 4686–4702 (2016).

23. Dousset, V. et al. Lysolecithin-induced demyelination in primates: preliminary in vivo study with MR and magnetization transfer. AJNR Am. J. Neuroradiol. 16, 225–31 (1995).

24. He, Y., Aznar, S., Siebner, H. R. & Dyrby, T. B. In vivo tensor-valued diffusion MRI of focal demyelination in white and deep grey matter of rodents. Neuroimage Clin. 30, 102675 (2021).

25. Lehto, L. J. et al. Lysophosphatidyl Choline Induced Demyelination in Rat Probed by Relaxation along a Fictitious Field in High Rank Rotating Frame. Front. Neurosci. 11, 433 (2017).

26. Jeffery, N. D. & Blakemore, W. F. Remyelination of mouse spinal cord axons demyelinated by local injection of lysolecithin. J. Neurocytol. 24, 775–781 (1995).

27. Raposo, C. & Schwartz, M. Glial scar and immune cell involvement in tissue remodeling and repair following acute CNS injuries. Glia 62, 1895–190. (2014).

28. West, K. L. et al. Myelin volume fraction imaging with MRI. Neuroimage 182, 511–521 (2018).

29. Kjer, H. M. et al. Bridging the 3D geometrical organisation of white matter pathways across anatomical length scales and species. Elife 13, (2025).

30. Berg, R. C. et al. Comparing myelin-sensitive magnetic resonance imaging measures and resulting g-ratios in healthy and multiple sclerosis brains. Neuroimage 264, 119750 (2022).

31. Graham, D. I. Triethyltin Sulfate-Induced Neuropathy in Rats. Arch. Neurol. 33, 40 (1976).

32. Brownstein, K. R. & Tarr, C. E. Spin-lattice relaxation in a system governed by diffusion. Journal of Magnetic Resonance (1969) 26, 17–24 (1977).

33. Chu, T. et al. Dynamic response of microglia/macrophage polarization following demyelination in mice. J. Neuroinflammation 16, 188 (2019).

34. Keough, M. B., Jensen, S. K. & Yong, V. W. Experimental Demyelination and Remyelination of Murine Spinal Cord by Focal Injection of Lysolecithin. Journal of Visualized Experiments 2015, 52679 (2015).

35. Hertanu, A. et al. T1D-weighted ihMT imaging – Part II. Investigating the long- and short-T1D components correlation with myelin content. Comparison with R1 and the macromolecular proton fraction. Magn. Reson. Med. 87, 2329–2346 (2022).

36. Dula, A. N., Gochberg, D. F., Valentine, H. L., Valentine, W. M. & Does, M. D. Multiexponential T2, magnetization transfer, and quantitative histology in white matter tracts of rat spinal cord. Magn. Reson. Med. 63, 902–909 (2010).

37. Harkins, K. D., Dula, A. N. & Does, M. D. Effect of intercompartmental water exchange on the apparent myelin water fraction in multiexponential T2 measurements of rat spinal cord. Magn. Reson. Med. 67, 793–800 (2012).

38. Helms, G., Dathe, H., Kallenberg, K. & Dechent, P. High-resolution maps of magnetization transfer with inherent correction for RF inhomogeneity and T1relaxation obtained from 3D FLASH MRI. Magn. Reson. Med. 60, 1396–1407 (2008).

39. Varma, G., Duhamel, G., de Bazelaire, C. & Alsop, D. C. Magnetization transfer from inhomogeneously broadened lines: A potential marker for myelin. Magn. Reson. Med. 73, 614–622 (2015).

40. Mezer, A. et al. Quantifying the local tissue volume and composition in individual brains with magnetic resonance imaging. Nat. Med. 19, 1667–1672 (2013).

41. Levesque, I. R. et al. Quantitative magnetization transfer and myelin water imaging of the evolution of acute multiple sclerosis lesions. Magn. Reson. Med. 63, 633–640 (2010).

42. Stikov, N. et al. In vivo histology of the myelin g-ratio with magnetic resonance imaging. Neuroimage 118, 397–405 (2015).

43. Mohammadi, S. & Callaghan, M. F. Towards in vivo g-ratio mapping using MRI: Unifying myelin and diffusion imaging. J. Neurosci. Methods 348, 108990 (2021).

44. Mohammadi, S. et al. Whole-Brain In-vivo Measurements of the Axonal G-Ratio in a Group of 37 Healthy Volunteers. Front. Neurosci. 9, 441 (2015).

45. Yarnykh, V. L. Fast macromolecular proton fraction mapping from a single off-resonance magnetization transfer measurement. Magn. Reson. Med. 68, 166–178 (2012).

46. Piredda, G. F. et al. Fast and high-resolution myelin water imaging: Accelerating multi-echo GRASE with CAIPIRINHA. Magn. Reson. Med. 85, 209–222 (2021).

47. De Groot, M. et al. Changes in normal-appearing white matter precede development of white matter lesions. Stroke 44, 1037–1042 (2013).

48. Hinman, J. D. & Abraham, C. R. What’s Behind the Decline? The Role of White Matter in Brain Aging. Neurochem. Res. 32, 2023–2031 (2007).

49. Pannese, E. Morphological changes in nerve cells during normal aging. Brain Struct. Funct. 216, 85–89 (2011).

50. Schmitt, A., Simons, M., Cantuti-Castelvetri, L. & Falkai, P. A new role for oligodendrocytes and myelination in schizophrenia and affective disorders? Eur. Arch. Psychiatry Clin. Neurosci. 269, 371–372 (2019).

51. Murayama, R., Cai, Y., Nakamura, H. & Hashimoto, K. Demyelination in psychiatric and neurological disorders: Mechanisms, clinical impact, and novel therapeutic strategies. Neurosci. Biobehav. Rev. 174, 106209 (2025).

52. Franklin, R. J. M., Bodini, B. & Goldman, S. A. Remyelination in the Central Nervous System. Cold Spring Harb. Perspect. Biol. 16, a041371 (2024).

53. Trujillo, P. et al. Contrast mechanisms associated with neuromelanin-MRI. Magn. Reson. Med. 78, 1790–1800 (2017).

54. Birkl, C. et al. The influence of brain iron on myelin water imaging. Neuroimage 199, 545–552 (2019).

55. Smith, K. J., Blakemore, W. F. & McDonald, W. I. The restoration of conduction by central remyelination. Brain 104, 383–404 (1981).

56. Klugmann, M. et al. Assembly of CNS Myelin in the Absence of Proteolipid Protein. Neuron 18, 59–70 (1997).

57. Paxinos, G. & Watson, C. The Rat Brain Atlas in Stereotaxic Coordinates. The Rat Brain in Stereotaxic Coordinates: 6th Edition (Academic Press, 2013).

58. Gochberg, D. F. & Gore, J. C. Quantitative magnetization transfer imaging via selective inversion recovery with short repetition times. Magn. Reson. Med. 57, 437–441 (2007).

59. Nyga rd, K. et al. ForMAX – a beamline for multiscale and multimodal structural characterization of hierarchical materials. J. Synchrotron Radiat. 31, 363–377 (2024).

60. Gu rsoy, D., De Carlo, F., Xiao, X. & Jacobsen, C. TomoPy: a framework for the analysis of synchrotron tomographic data. J. Synchrotron Radiat. 21, 1188–1193 (2014).

61. Riise, J. & Pakkenberg, B. Stereological estimation of the total number of myelinated callosal fibers in human subjects. J. Anat. 218, 277–28. (2011).

62. Cardiff, R. D., Miller, C. H. & Munn, R. J. Manual Hematoxylin and Eosin Staining of Mouse Tissue Sections. Cold Spring Harb. Protoc. 2014, 655–658 (2014).

63. Georgiadis, M. et al. Nanostructure-specific X-ray tomography reveals myelin levels, integrity and axon orientations in mouse and human nervous tissue. Nat. Commun. 12, 2941 (2021).

64. Virtanen, P. et al. SciPy 1.0: fundamental algorithms for scientific computing in Python. Nat. Methods 17, 261–272 (2020).

65. Seabold, S. & Perktold, J. Statsmodels: Econometric and Statistical Modeling with Python. In Proceedings of the 9th Python in Science Conference vol. 57 92–96 (2010).

66. Terpilowski, M. scikit-posthocs: Pairwise multiple comparison tests in Python. J. Open Source Softw. 4, 1169 (2019).

67. Klein, S., Staring, M., Murphy, K., Viergever, M. A. & Pluim, J. elastix: A Toolbox for Intensity-Based Medical Image Registration. IEEE Trans. Med. Imaging 29, 196–205 (2010).

68. Lakens, D. Equivalence Tests: A Practical Primer for t Tests, Correlations, and Meta-Analyses. Soc. Psychol. Personal. Sci. 8, 355–362 (2017).

