## Supplementary Information for "Quantitative MRI-based assessment of the ultrastructural compactness of myelin lamellae"

### Supplementary Figure 1. Analysis of longitudinal changes during postmortem degeneration.

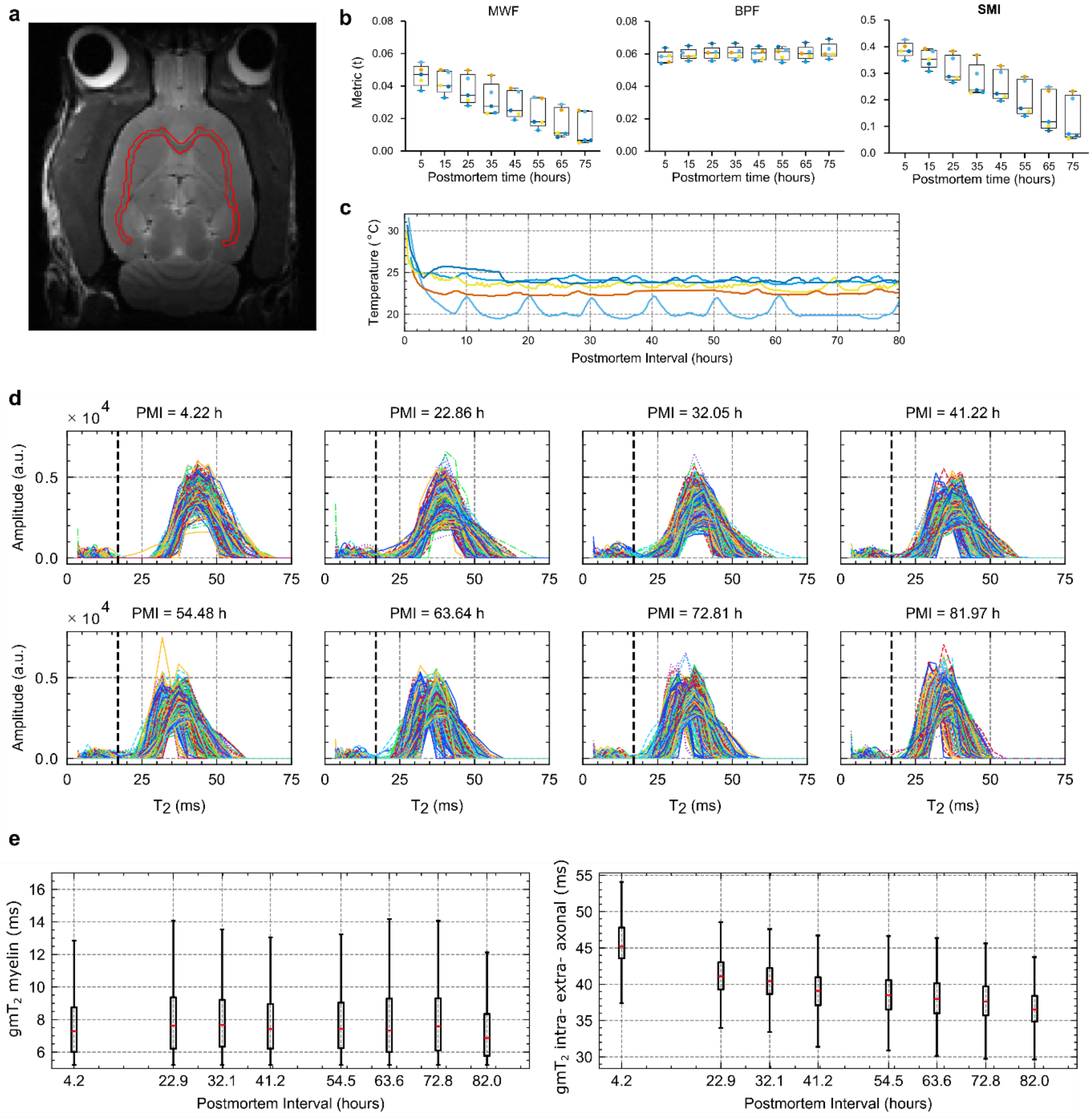


**Supplementary Figure 1. Analysis of longitudinal changes during postmortem degeneration. a.** Structural image with the white matter region of interest delineated in red for a representative animal. **b.** Longitudinal evolution of the interpolated qMRI metrics in eight time-points for statistical analysis. **c.** Body temperature trends for each animal. **d.** Representative $T_2$ spectra in the region of interest in each acquisition block for one animal. The dashed black line indicates the cut-off value for MWF quantification. **e.** Boxplots for the voxel-wise geometrical mean $T_2$ values of the myelin compartment (left) and the intra-extra axonal compartment (right) in the region of interest for one animal.

### 1 Supplementary Note 1: Prospective power analysis for the equivalence and stabilization tests in the LPC study.

The equivalence tests reported in the LPC-induced demyelination results section did not establish equivalence to naive baseline or stabilization between 21 and 28 dpi for any metric. Because a non-significant equivalence test can reflect either an actual non-negligible difference or insufficient statistical power, we performed a prospective power analysis based on the pilot data reported here to estimate the cohort sizes that would be required for these same tests to reach conventional power (80%) at the pre-specified equivalence margin. Moreover, we additionally examined how MWF, BPF and SMI covaried across animals at late recovery stages to support the interpretation of the results. This analysis is reported here as supplementary context for interpreting the null equivalence results.

#### 1.1 Methods

**Power analysis**

Required sample sizes were estimated analytically using the Welch-corrected two one-sided test (TOST) framework with Satterthwaite approximate degrees of freedom, applied to the mean difference with a normal-theory confidence interval[1]. We applied Welch's correction to avoid the assumption of equal variances across the groups being compared. The equivalence margin was defined as half the standard deviation of the naive reference distribution for each metric, matching the margins in the main equivalence tests. Per-group means and standard deviations were estimated from the pilot data (n=5 per group) and were kept separate per group. Given the limited sample size of mean and standard deviation estimates (n=5), required sample sizes should be interpreted as order-of-magnitude guidance rather than precise targets.

**Between animal covariation across metrics**

Spearman rank correlations were computed between the lesion-to-contralateral ratios of MWF, BPF and SMI across animals at 21 and 28 dpi (pooled, n = 10). Because both MWF and BPF ratios increase between 21 and 28 dpi, the correlation on pooled data could partially reflect the recovery time trend rather than the isolated correlation between metrics across animals. Thus, correlations were computed after centring each metric on the median of its own timepoint group. The correlation between MWF and BPF was the primary test of interest; correlations with SMI are reported descriptively. Similar to the power analysis section, with n=10, the correlations found here should be considered suggestive rather than conclusive.

#### 1.2 Results

Required sample sizes per group to reach 80% power under the assumption that the groups are equal (i.e., the true difference is 0) are reported in Supplementary Table 1.

**Supplementary Table 1:** Prospective power analysis sample requirements for assuming true no differences with respect to naive or between the 21 and 28 dpi groups.

| Analysis | Metric | Comparison | Required *n* per group |
|---|---|---|---|
| Equivalence vs. naive | MWF | naive vs. 21 dpi | 44 |
| | MWF | naive vs. 28 dpi | 226 |
| | BPF | naive vs. 21 dpi | 316 |
| | BPF | naive vs. 28 dpi | 826 |
| | SMI | naive vs. 21 dpi | 43 |
| | SMI | naive vs. 28 dpi | 41 |
| Stabilization | MWF | 28 vs. 21 dpi | 208 |
| | BPF | 28 vs. 21 dpi | 1126 |
| | SMI | 28 vs. 21 dpi | 33 |

We additionally found that MWF and BPF ratios were positively correlated across animals at late recovery timepoints ($\rho = 0.63$, $p = .052$). In contrast, SMI was not significantly correlated with MWF ($\rho = 0.20$, $p = .58$) or BPF ($\rho = -0.34$, $p = .33$).

#### 1.3 Interpretation

Assuming no differences, establishing statistical equivalence to the naive distribution would require from 41 to 826 animals per group, and showing stabilization would require between 33 to 1126 animals (Supplementary Table 1), cohort sizes unrealistic under this study design. Since the equivalence margins for each metric were defined from the variability of its corresponding naive reference distribution, the differences between cohort sizes mainly reflect the between-animal variability within lesion groups compared to the naive tissue. For example, BPF's equivalence margin is narrow (±0.007) because the naive cohort is homogeneous, while the between-animal spread in the lesion cohort is relatively large. For MWF, required cohort sizes increased substantially from 21 to 28 dpi, reflecting greater between-animal variability at the later timepoint. Since we found that MWF and BPF ratios were also positively correlated across animals at late recovery stages, this suggests that part of their variability is shared. Such shared variation could arise from differences between animals in the recovery trend, the initial lesion extent, or the overlap between lesion and region of interest.

In contrast, required cohort sizes for SMI remained relatively small and stable (33–43 animals per group) for both analyses. This is consistent with the hypothesis that SMI reflects the organisation of myelin rather than its amount: shared variation by MWF and BPF is expected to cancel in SMI, and the remaining variability would be low if the compactness of remyelinated sheaths is relatively homogeneous across animals at late recovery stages.

Overall, these results offer a reference for prospective power calculations and indicate that SMI is the most feasible metric for stabilization testing. Future studies can reduce sample size requirements by using a longitudinal design that images the same animals at 21 and 28 dpi, creating a practical approach to establishing equivalence or stabilization.

### Supplementary references

1. Lakens, D. Equivalence Tests: A Practical Primer for t Tests, Correlations, and Meta-Analyses. *Soc. Psychol. Personal. Sci.* **8**, 355–362 (2017).
